# Cued Naming and the left inferior frontal cortex

**DOI:** 10.1101/2021.02.10.430676

**Authors:** Rachel Holland, Jennifer T. Crinion

**Affiliations:** Institute of Cognitive Neuroscience, University College London, 17 Queen Square, London, WC1N 3AR, UK; Division of Language and Communication Science, City University London, London, EC1V 0HB, UK

**Author notes:** Correspondence should be addressed to: Professor Jenny Crinion.

## Abstract

Clinical studies have shown that naming can be behaviorally facilitated by priming, e.g., phonemic cues reduce anomia. Rehabilitation of language is argued to rely upon the same processes of priming in healthy speakers. Here we show, in healthy older adults, the immediate facilitatory behavioral and neural priming elicited by phonemic cues presented during an fMRI experiment of overt naming; thus, bridging the gap between lesion and neuroimaging studies. Four types of auditory cues were presented concurrently with an object picture (e.g., cat): (i) word (i.e., the target name (/kat/), (ii) initial phoneme segment (e.g., /ka/), (iii) final phoneme segment (/at/), or (iv) acoustic (noise) control cue. Naming was significantly faster with word, initial and final phonemic cues compared to noise; and word and initial cues compared to final cues, with no difference between word and initial cues. A neural priming effect – a significant decrease in neural activity – was observed in the left inferior frontal cortex (LIFC, pars triangularis, BA45) and the anterior insula bilaterally consistent with theories of primed articulatory encoding and post-lexical selection. The reverse contrast revealed increased activation in left posterior dorsal supramarginal gyrus for word cues that, we argue, may reflect integration of semantic and phonology processing during word rather than phonemic conditions. Taken together, these data from unimpaired speakers identified nodes within the naming network affected by phonemic cues. Activity within these regions may act as a possible biomarker to index anomic individuals’ responsiveness to phonemically cued anomia treatment.

## Introduction

Naming an object is a complex task that calls on several different processing stages that precede articulation, most importantly conceptual recognition and retrieval of its spoken name (phonemic form) (Blackford et al., 2012, Levelt et al., 1991). The inability to efficiently name common everyday objects (anomia) is a hallmark symptom of aphasia, regardless of severity (Crinion and Leff, 2007, Goodglass, 1993). Importantly, naming performance can be improved in some aphasic individuals by repeatedly pairing a picture of the to-be-named object with an auditory cue that carries some speech sound (phonemic) information related to the object in question (Kendall et al., 2008, Pease and Goodglass, 1978). It has been argued that successful phonemic cuing therapy relies upon the same processes that underlie priming in unimpaired speakers (Best et al., 2002; Nickels 2002). That is, phonemic cues prime retrieval of the correct phonological form (Miceli et al., 1996). Indeed, naming can be primed in unimpaired speakers when the target picture (e.g. of a cat) is presented with auditory phonemic whole word cues (e.g., /kat/) or part segment cues (e.g., /ka/) (Starreveld, 2000). Yet, despite the longstanding use of phonemic cues clinically and the assertion that treatment-induced naming improvements rely upon a normal priming network, the neural underpinnings of phonemically-cued priming of naming in unimpaired speakers remains unclear. Here, we explore how cues presented simultaneously with a picture of a to-be-named object, prime the behavioral and neural naming responses in unimpaired older adult speakers using functional magnetic resonance imaging (fMRI).

Retrieval of the correct phonological form from semantics – a key processing step in object naming – has been associated with enhanced activation in the left frontal cortex (Price, 1998). The inferior aspect of frontal cortex (Mechelli et al., 2007) and the insula cortex (Dronkers, 1996; Wise et al., 1999) have also been implicated in the later stages of speech production, specifically generation of the appropriate articulatory code and mapping heard speech sounds onto their articulatory codes (Elmer et al., 2011). Given this association, we predicted a significant reduction in, or priming of the neural response in left inferior frontal cortex (LIFC) and bilateral insula as articulatory processing demands are reduced by the provision of auditory cues during overt naming.

To date, only two neuroimaging studies have explored the priming effect of auditorily presented whole words that shared an initial phoneme with the target name (Abel et al., 2009, Abel et al., 2012, de Zubicaray and McMahon, 2009). That is, presenting the whole word cue “hat” primes naming a picture of a hand. While these latter studies address priming by phonologically related whole words, the auditory cues differ widely from those used in naming therapy. Here we adopted a cuing task that closely mirrors the type of cues used clinically to identify where within the naming network phonemic cues elicit priming in unimpaired individuals, thus, bridging the gap between lesion studies and functional neuroimaging studies of language production in healthy speakers.

## Materials and Methods

### Participants

22 healthy right-handed native speakers of English (14 females, mean age 66 years; range 49-88 years) participated in the study. All had normal hearing and no previous history of neurological or psychiatric disease or metallic implants. All participants gave their written and informed consent to participate in the study and the study was approved by the Central London Research Ethics Committee.

### Stimuli

The pictorial stimulus set consisted of 107 black and white line drawings of objects drawn from those of Szekely et al. (2004). All object names were monosyllabic and consonant-vowel-consonant (CVC) in terms of phonological structure. For example, the word /kat/ is translated as [kæt] in the International Phonetic Alphabet and has a phonological structure of CVC. The word /fish/, although consisting of CVCC in its orthographic form, also translates to CVC in terms of its phonological form (i.e., [fI∫]). Using the norms (Szekely et al., 2004) from the International Picture Naming Project (IPNP) all selected objects had a name agreement (i.e., percentage of participants producing the target name) of greater than 75%.

Each picture was presented simultaneously with an auditory cue. Auditory cues were either: (i) a whole word cue (i.e., the target name (cat: /kat/), (ii) an initial phoneme segment (e.g., /ka/), (iii) a final phoneme segment (/at/), or an unintelligible auditory control (noise) cue. To generate the auditory cues, each target name for each target object was digitally recorded (sampling rate 44.1KHz) from a male native speaker of English in a soundproof room. For word cues, the auditory cue was the spoken whole word token. To generate the initial and final phoneme cues each whole word token was cropped at either the (i) offset of the vowel to form the initial cue (e.g., /ka/) or (ii) the onset of the vowel to form the final cue (e.g., /at/). Initial and final cues were then matched for total auditory duration. To generate a control cue for each target object, each of the spoken word cues that accompanied each target picture were spectrally rotated (Blesser, 1972) and then submitted to a noise-vocoding routine (Shannon et al., 1995) using a single level of filter band noise vocoding. This procedure leaves the temporal envelope of the spoken token unaltered and preserves the spectro-temporal complexity, whilst rendering the auditory signal unintelligible by inverting the frequency spectrum. This control condition has successfully been used in previous imaging studies (Narain et al., 2003, Obleser et al., 2007, Scott et al., 2000).

### Procedure

Four functional runs were acquired within one scanning session. Each of the 107 picture stimuli was presented only once during a functional run with one of the four different cue types. Over the scanning session, therefore, each picture was seen a total of four times – presented once with each auditory cue (word, initial, final phoneme segment, or a noise control). Within each run, similar numbers of each of the four cue types were presented. The order of pictures and accompanying cues was pseudo-randomized (more than three trials with the same cue type were avoided) and order of presentation was counterbalanced both within and across participants.

Each picture was displayed for 2500 ms and preceded by a 1000 ms fixation cross. Auditory cues were presented simultaneously with each picture (stimulus onset asynchrony SOA = 0 ms). The time at which the phonemic cue is delivered has important consequences for priming. Previous studies have found robust priming effects with a SOA of 0 ms (de Zubicaray and McMahon, 2009, Meyer and Schriefers, 1991, Schriefers et al., 1990, Starreveld, 2000), with phonological priming effects diminishing with early SOAs (c.f., Abel et al., 2009). Trials were presented in short blocks of 6 stimuli, separated by a fixation-only rest period of 7 seconds. The inter-trial interval was set to 3920 ms so as to jitter the onset on each trial across acquired volumes in order to vary the spatial acquisition of data.

Participants were instructed to name the picture as quickly and as accurately as possible. Overt spoken responses were recorded in the scanner using a dual-channel, noise-cancelling fiber optical microphone system (FOMRI III http://www.optoacoustics.com). Each response was reviewed offline to verify manual recording of accuracy and used to determine trial-specific reaction times for each participant. Auditory cues were delivered via an MR-compatible set of headphones (MR Confon, Magdeburg, Germany; www.mr-confon.de).

### Image acquisition

Whole-brain imaging was performed on a 3T Siemens TIM-Trio system (Siemens, Erlangen, Germany) at the Wellcome Trust Centre for Neuroimaging. Using a 12-channel head coil T2*-weighted echo-planar images (EPI) with BOLD contrast were acquired. Each EPI comprised 48 AC/PC-aligned axial slices with sequential ascending acquisition; slice thickness of 2 mm, 1 mm inter-slice gap and a 3 x 3 mm in-plane resolution. Volumes were acquired with a repetition time (TR) of 3360 ms per volume and the first six volumes of each session were discarded to allow for T1 equilibration effects. A total of 180 volume images (174 volumes of interest, 6 dummy scans) were acquired in four consecutive runs, each lasting approximately 10 minutes. Prior to the first functional run of each scanning session, a gradient field map was acquired for each participant for later B0 field distortion correction of functional images. The same scanner and hardware were used for the acquisition of all images.

The functional data were preprocessed using Statistical Parametric Mapping software (SPM8; www.fil.ion.ucl.ac.uk/spm) running under Matlab 2008b (MathWorks, Natick, MA). All volumes of interest from each participant were realigned and unwarped, using session and subject-specific voxel displacement maps (Hutton et al., 2002). The functional images were then spatially normalized to the standard T2* template within SPM normalization software. Functional data were spatially smoothed, with a 8mm full-width at half-maximum isotropic Gaussian kernel to allow for residual variability after spatial normalization and to permit application of Gaussian random field theory for corrected statistical inference.

Statistical analyses were first performed in a subject-specific fashion. To remove low-frequency drifts, the data were high-pass filtered using a set of discrete cosine functions with a cut-off period of 128 sec. Each cue type was modelled separately as an event by convolving it with the SPM canonical haemodynamic response function (HRF). We used presentation of the pictorial and auditory cue as the onset of the event to model the preparatory naming response. Trial-specific reaction times were also included in the regression analysis to model speech onset time. Movement realignment parameters were included as covariates of no interest. The resulting stimulus-specific parameter estimates were calculated for all brain voxels using the General Linear Model. Contrast images were computed for (i) each cue type relative to rest and (ii) overall naming reaction time for whole brain analyses at the second level. The statistical threshold was set to p<0.05 after family wise error (FWE) correction for multiple comparisons across the whole brain.

## Results

### Behavioral data

Naming responses remained accurate throughout (Table 1). Trials in which participants gave an incorrect naming response were excluded from the analysis (1.5% of trials in total). Data were analyzed by subjects (F_1_) and by items (F_2_). A repeated measures ANOVA was conducted on the reaction time means with cue type as a within subject variable. The results indicated a significant main effect of cue type by subjects and by items (F_1_(3,63) = 68.0, *p* < 0.001; see Figure 1; F_2_(3,318) =101.19, p<0.001). Planned post hoc comparisons showed that, by subjects and by items, naming responses were significantly primed when pictures were accompanied by phonemic cues (word, initial and final phoneme segment) compared to the control cue condition (all phonemic cue condition comparisons relative to control cue: p-values < 0.001, Figure 1).

**Table 1:** Mean correct reaction time (RT) with standard error of the mean (SEM) in parentheses and proportion accuracy. (ms) = milliseconds. Accuracy expressed as a proportion and calculated for each cue condition by dividing the total number of correct naming responses by the total number of items presented within the condition.

| Cue condition | Response time (ms) | Accuracy (prop.) |
| --- | --- | --- |
| Word | 842 (27) | 0.998 |
| Initial | 841 (24) | 0.992 |
| Final | 934 (21) | 0.989 |
| Control | 984 (24) | 0.964 |

**Figure 1:**
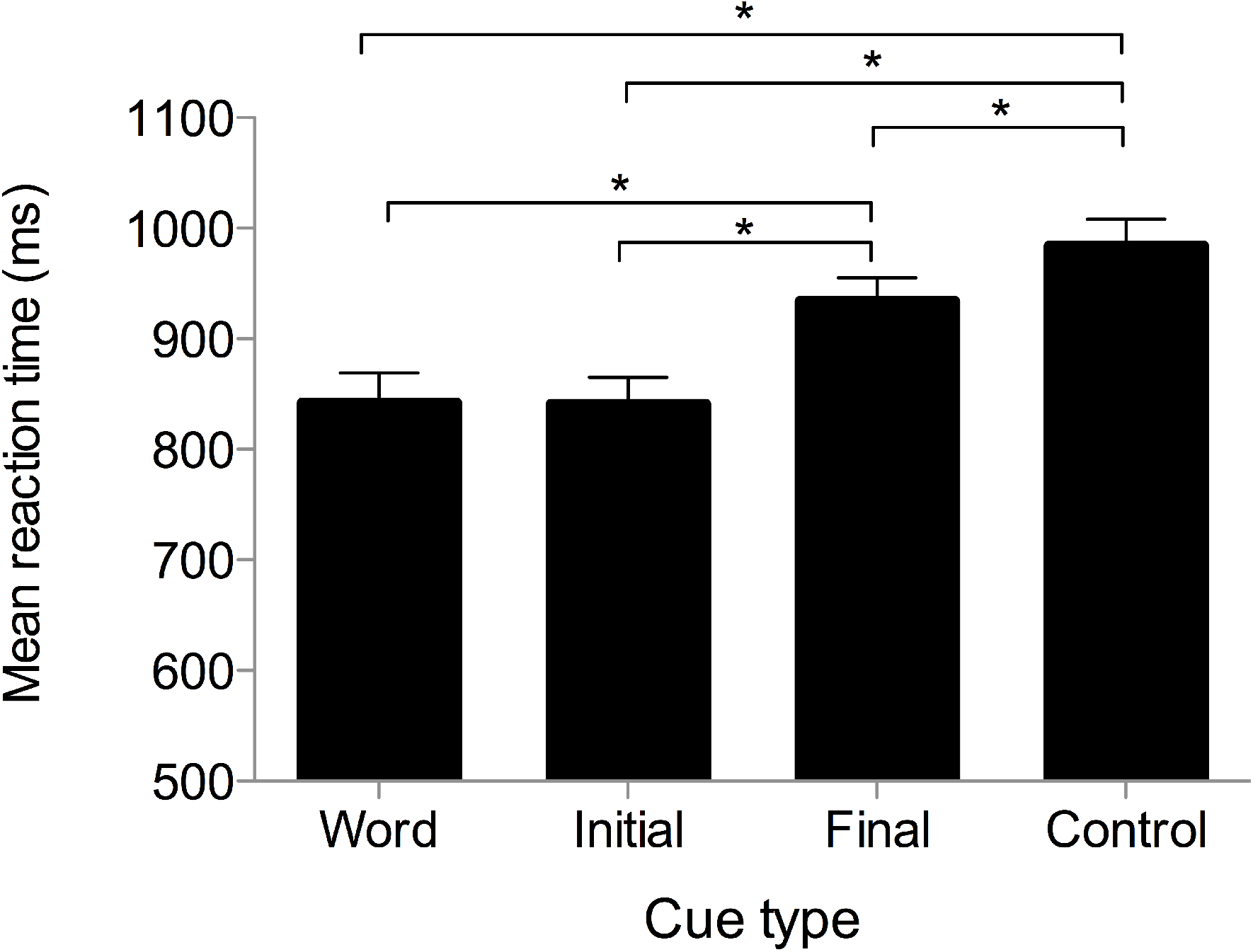
Mean reaction time (RT) by subjects and standard error of mean (+SEM) for each cue type during overt fMRI naming (*N* = 22). * indicates a significant difference at P < 0.001. (ms) = milliseconds.

Comparisons between cue type revealed that, by subjects, both word and initial phoneme segment cues facilitated naming responses more than final phoneme segment cues (both cue types compared to final: p < 0.001), with no significant difference between the word and initial phoneme segment cues (t(21) = p = −0.08, p = 0.94). By-item analysis revealed significant differences between all cue types, including word and initial phoneme segment cues (all p-values <0.001).

### fMRI results

#### Naming network

Overall, overt object naming irrespective of cue type activated an extensive bilateral fronto-temporal neural network relative to rest blocks. Peak activation associated with overt naming (summed over cue type) relative to rest was observed bilaterally in the superior temporal gyrus extending to the precentral gryri and middle cingulate cortex. A distinct cluster of activation was identified in the left insula, the right temporal and middle occipital gyrus extending to the inferior aspect of the lobe. Distinct peaks within the cerebellum and calcarine sulcus were also identified bilaterally (Figure 2a).

**Figure 2:**
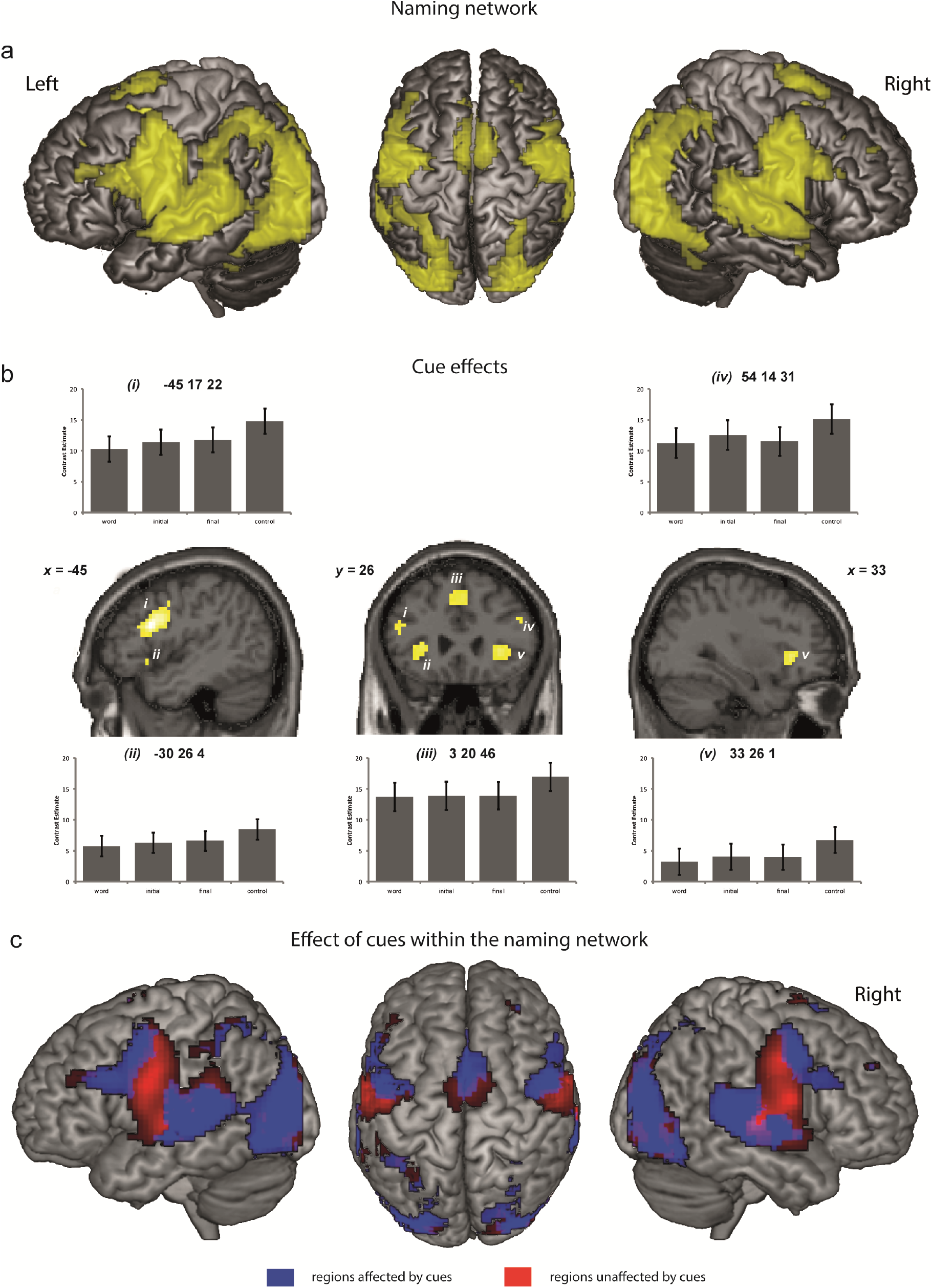
Neural response, overlaid on a canonical brain, for contrasts of interest: (**a**) the simple main effect of naming (summed over cue type) relative to rest; (**b**) the facilitatory effect of speech cues (word, initial and final phoneme segment) relative to control with mean beta values for each peak voxel plotted in graphs *i* – *v.* Plots show decreased BOLD response associated with speech cues compared to control in (*i*) left IFG (−45 17 22), (*ii*) left insula (−30 26 4), (*iii*) right middle cingulate cortex (3 20 46), (*iv*) right IFG (54 14 31) and (*v*) right insula; and (**c**) the regions within the naming network that are and are not affected by speech cues. Activation illustrated in panels a,b and c was corrected in height for multiple comparisons across the whole brain (FWE; P < 0.05). All coordinates are in MNI space (x,y,z). IFG = inferior frontal gyrus.

#### Cue effects

##### (i) Simple main effect of cues versus control (contrast: control > cues)

Neural responses mirrored behavior: we identified a significant priming of the BOLD response associated with phonemic cues relative to the control cue (corrected for whole brain comparisons (FWE), P<0.05) in the left inferior frontal gyrus (IFG; pars triangularis, x= −45, y= 17, z= 22, BA45; Z score = 6.87) with a distinct sub-peak of the cluster in left precentral gyrus (−42 8 31; Z score = 6.32).

Separate peaks in the insula were identified bilaterally (left: x=-30, y= 26, z= 4; Z score = 5.44; right: x= 33, y= 26, z= 1; Z score = 5.69) extending anteriorly to the pars orbitalis (x=-45, y=20, z=-8). Two further distinct peaks were identified in the right inferior frontal gyrus (pars opercularis: x=54, y=14, z=31; Z score =5.57) and the right posterior medial frontal region – specifically the right middle cingulate (x=3, y=20, z=46; Z score = 5.46) with sub-peaks extending to the left superior middle gyrus (x=-35, y=35, z=37; Z score = 4.61). See Figure 2b; Table 2.

**Table 2:**
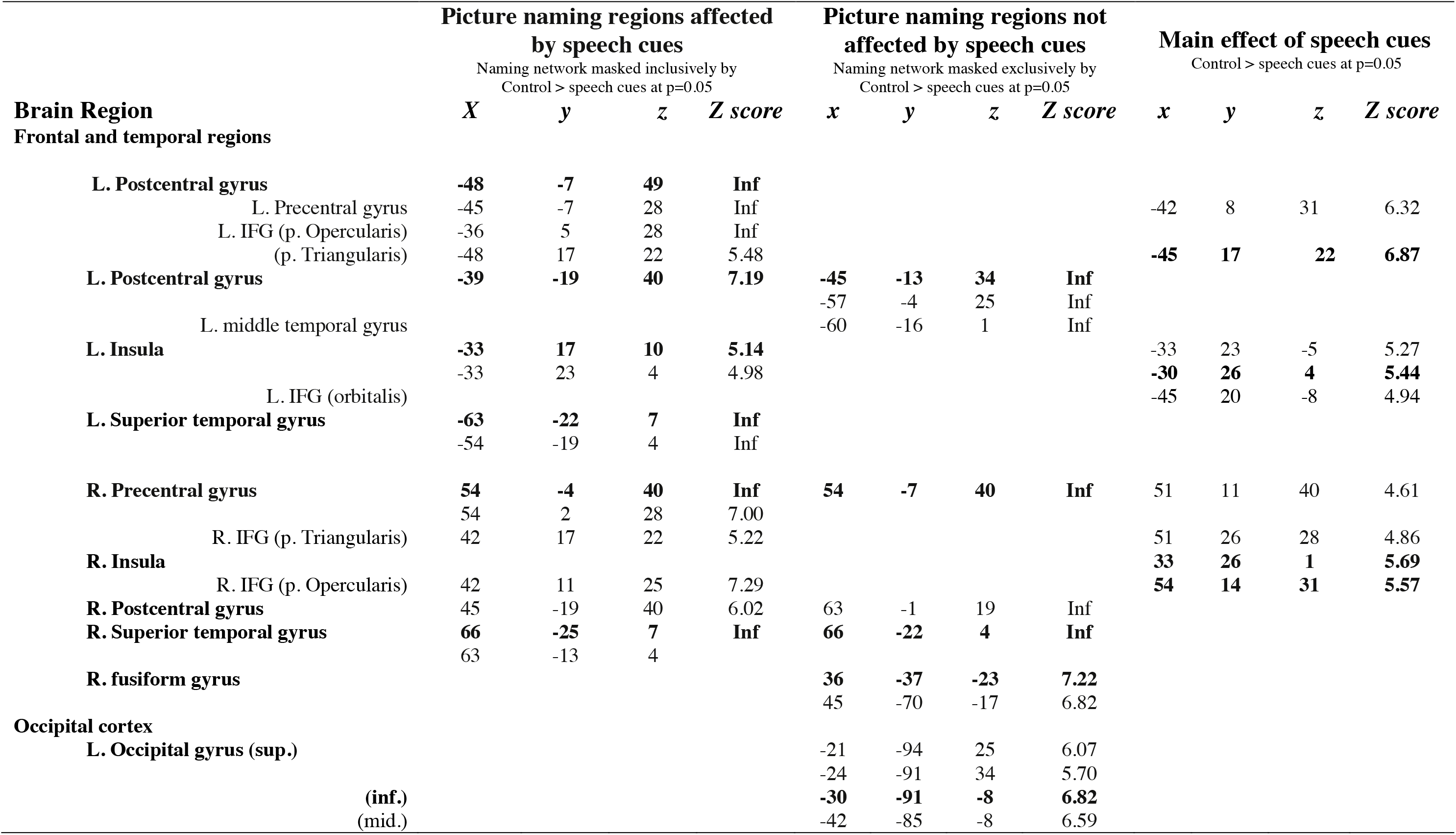
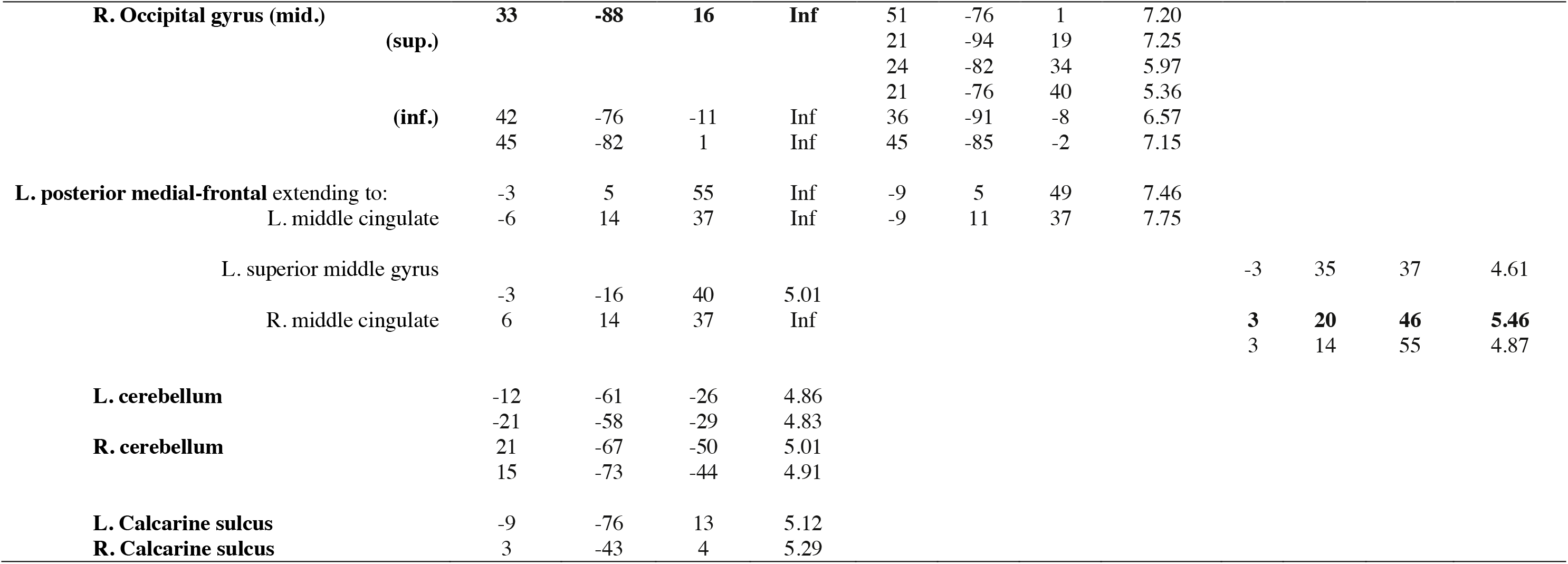
Brain regions significantly activated for contrasts of interest. Peak activation in brain regions that show a facilitatory effect of speech cues (word, initial, final phoneme segment) relative to control cue are reported in the far right column. The differential effect of cues on activation within the picture naming regions is also reported. For large clusters, the peak voxels within a cluster are labelled in bold typeface and prominent local maxima are listed separately (indented). All coordinates are in MNI space (x,y,z) and peak voxel Z-scores reported. MNI coordinate labels determined using Eickhoff et al., (2005) SPM Anatomy Toolbox used for labelling. Peak-labels that are consistent across contrasts are reported on the same line with corresponding MNI coordinates listed.

##### (ii) Picture naming regions affected by cues

To determine which picture naming regions were affected by cues additional analyses were conducted (contrast: naming network masked inclusively by cue effects at p=0.05). Results indicated that regions within the naming network that were affected by cues included the postcentral gyri extending anteriorly to the precentral gyrus and encompassing the pars opercularis and pars triangularis. A peak within the left postcentral gyrus was also identified which extended into the middle temporal gyrus. Distinct peaks were identified in the bilateral insula extending to the pars orbitalis on the left and the pars opercularis on the right; the superior temporal gyrus. On the right, a distinct cluster of activation was identified in the right fusiform gyrus. Peaks of bilateral activation were also seen within the occipital gyrus, with sub-peaks identified in the superior, middle and inferior gyri of both hemispheres. Significant peaks of activation within the midline were also noted. A distinct cluster in the left posterior medial frontal region was identified with specific sub-peaks in bilateral middle cingulate and left superior middle gyrus.

##### (iii) Picture naming regions not affected by cues

To determine which picture naming regions were not affected by cues the activation associated with the naming network was masked exclusively by cue effects at p=0.05. Results identified regions within the naming network that did not partner with the simple effects of the phonemic cue (see Table X). These regions included peaks within the left postcentral gyrus, the right precentral gyrus – despite the cluster from simple cue effects extending to these regions, the maximal effect of the cue was located in the RIFG – and right fusiform and occipital regions. Despite these regions being implicated in the naming network, they did not show a reliable effect of phonemic cues.

##### (iv) Within-cue effects

We also explored differential effects within cues (i.e., initial and final cues compared to word cues). Although word and initial cues elicited equivalent behavioral responses, it could be argued that differences between word and word-initial segment cues would be expected at a cognitive level. Word-cued trials meant that to name the item, subjects need only to repeat what they hear, whilst initial and final segment cues may require additional cognitive processes of lexical retrieval or articulatory planning. As can been seen from the plotted data for the cue effects, word cues consistently appear to elicit a greater degree of neural priming than initial and final phoneme segment cues (Figure 2b).

However, direct comparison of the neural response for word cues relative to initial phoneme segment cues indicated no reliable differences, which mirrored the behavior. In contrast, word versus final phoneme segment cues showed a greater priming of the neural response for word cues in the LIFC (x = − 42, y = 23, z = 19; Z = 3.63), but only at an uncorrected level. The reverse contrast of enhancement for word cues relative to initial and final cues revealed an increase in activation in left posterior cortex in inferior parietal cortex for cues (IPC: x = −57, y = −52, z = 43; Z score = 4.90, FWE corrected). Direct comparison of word versus initial and word versus final phoneme segment cues activated the corresponding cluster (IPC: x = −51, y = −55, z = 46; Z score = 3.91 and 3.89 respectively, see Figure 3).

**Figure 3:**
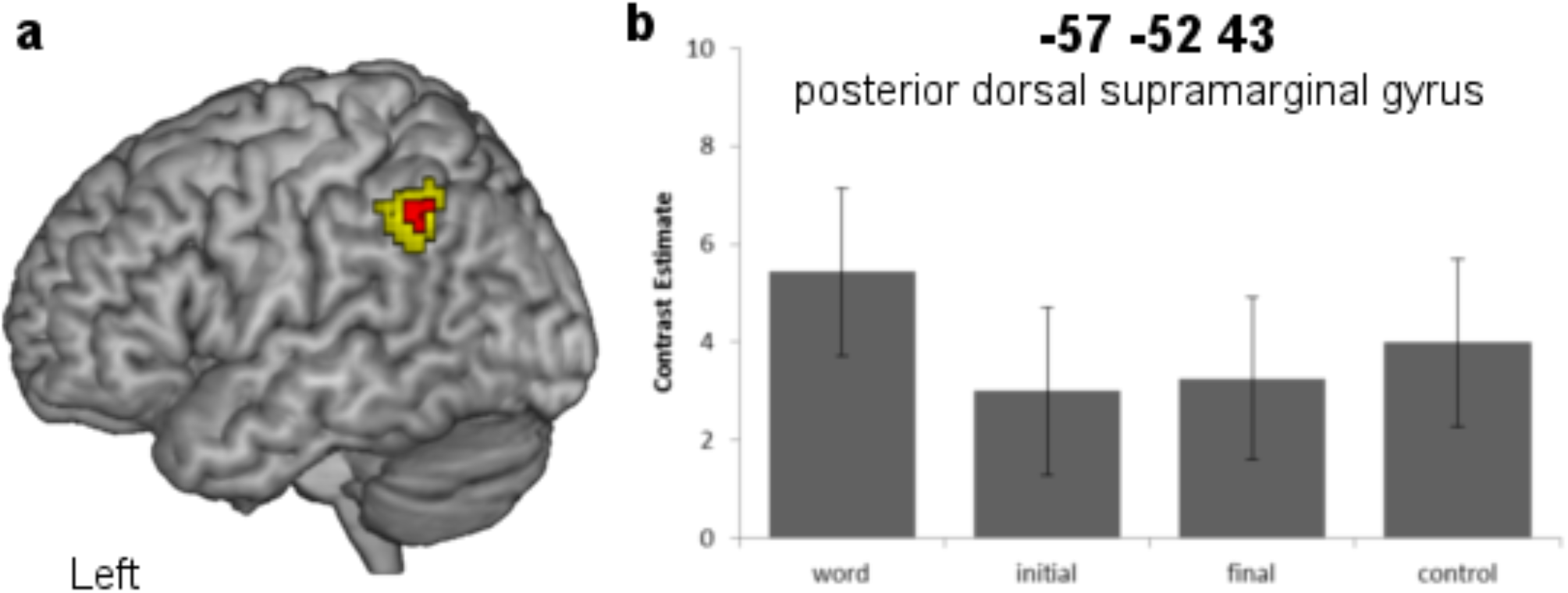
(**a**) Increased activation within the naming network for word cues relative to initial and final segment cue conditions. Contrast: word > initial + final masked inclusively by naming. Yellow cluster showing extent of activation at p=0.001 uncorrected and red cluster showing activation corrected for multiple comparisons across the whole brain (P=0.05) (**b**) mean beta value with SEM of each condition at peak voxel in left posterior dorsal supramarginal gyrus. All coordinates are in MNI space (x,y,z).

## Discussion

The inability to efficiently name a common object is a hallmark symptom of aphasia (Goodglass, 1993), which can be alleviated in some aphasic individuals by repeatedly pairing a picture of the to-be-named object with an auditory cue that carries some speech sound (phonemic) information related to the object in question (Kendall et al., 2008, Pease and Goodglass, 1978). Treatment-induced improvement in naming performance in some anomic individuals is argued to be akin to priming effects observed in unimpaired speakers (Best et al., 2002; Nickels 2002). Yet, the neural basis of phonemic priming of naming in unimpaired speakers remains unclear. During fMRI, we presented either whole word cues that were identical to the target name, or part words that shared either word-initial or -final phonemic segments to explore how phonemic cues affected the behavioral and neural naming response in unimpaired older adult speakers.

As predicted, all phonemic cues significantly primed naming response times compared to a control cue. Specifically, word and initial cues primed naming response time significantly more than final cues, with no reliable difference between word and initial cues. The magnitude of this behavioral priming effect was notably large and robust across participants (mean priming response relative to control cue: 111.67 ms) compared to other phonemic priming studies (c.f., Starreveld, 2000: range of mean phonemic priming effect: 38 - 92 ms). Auditory phonemic cues, therefore, can significantly prime the response to object names with high frequency and imageability, in an unimpaired language system indicating residual capacity within the naming network.

Importantly, these behavioral priming effects were mirrored by corresponding reductions in blood-oxygen-level dependent (BOLD) signal in the pars triangularis region (BA45) of the LIFC and the anterior insula bilaterally, as predicted. This pattern of response is consistent with the region being integral to the retrieval of the correct phonological form from semantics and generation of the appropriate articulatory code ready for overt speech (Elmer et al., 2011; Price, 2012; Wise et al., 1999).

Speech production is argued to be a naturally competitive process with word retrieval from semantics affected by the availability of competing alternatives. Neuroimaging studies reported increased activation in the LIFC with increased numbers of competing spoken responses (Kan and Thompson-Schill, 2004, Schnur et al., 2009). The provision of a phonemic cue, in contrast, reduces the number of competing responses by providing part of the phonological form that needs to be retrieved prior to articulation.

In addition to the neural priming of activation in LIFC, a reduction in BOLD signal was also observed in the insula bilaterally, as predicted. The bilateral insula have been associated with the later stages of speech production, highlighting its role in generation of the appropriate phonological or articulatory code (Mechelli et al., 2007). Specifically, the study reported increased activation in the insula bilaterally when two phonologically related words are produced sequentially. This profile of response was argued to reflect increased phonological processing demands associated with discriminating between two similar phonological or articulatory codes. In the present study, the primed neural response in the anterior insula bilaterally was significantly reduced by the provision of a phonemic cue, which is consistent with an explanation in terms of reduced phonological or articulatory retrieval demands. Indeed, the left anterior insula has been associated with retrieval of appropriate motor articulatory programs (Wise et al., 1999) and reported to be damaged in patients with apraxia of speech, which is a disruption to the motor articulatory planning of speech rather than production of speech sounds and articulatory movements *per se* (Dronkers and Ogar, 2004, Dronkers, 1996, however see Ackermann and Riecker, 2010, for a review). Furthermore, this region has shown hypometabolism (Nestor et al., 2003) and atrophy in patients with progressive non-fluent aphasia, a disorder characterized by non-fluent speech in the context of preserved comprehension (Gorno-Tempini et al., 2004). The right insula has been implicated in prosodic aspects of speech (Ackermann and Riecker, 2004, Bohland and Guenther, 2006) and spatiotemporal processing and coordination of speech (Geiser et al., 2008, Riecker et al., 2000).

The behavioral profile mirrored at the neural level as a facilitation of response in LIFC and bilateral insula is clear. However, it could be argued that a difference between word and phonemic segment cues would be expected at a cognitive and therefore neural level. That is, word-cued naming may simply reflect repetition. At a cognitive level, significantly greater priming of the neural response may be expected for word-cued items as the auditory cue completely overlaps with the phonological features of the target name, eliminating any selection demand. The reverse contrast, however, revealed that word cues preferentially engaged left inferior parietal cortex (IPC: x = −58, y = −54, z = 40). This enhanced activity associated with word cues was of particular interest. Its anatomical location in the posterior dorsal supramarginal gyrus (pdSMG) on the border between the angular gyrus associated with semantic processing and the anterior supramarginal gyrus associated with phonological processing suggests that the enhanced activation may be related to the integration of semantic and phonology processing (x = −44, y = −54, z = 46; Lee et al., 2007). This study is highly pertinent to the current result as not only did white matter tractography highlight the connection of this region to the anterior supramarginal gyrus and angular gyrus, but also activity within the region was positively correlated with the extent of spoken word vocabulary. Mechelli et al., (2007) also reported activation in an highly proximate area (x = −58, y = −52, z = 40) during overt production of semantic relative to phonological related word pairs further suggesting its role in aspects of semantic processing.

In summary, it has been argued that successful phonemic cuing therapy relies upon the same processes that underlie priming in unimpaired speakers. Here, we show for the first time, that phonemic cues that share only phonemic features with the target name, significantly prime behavior and the neural response in LIFC and bilateral insula. We propose this priming effect is consistent with theories of selection among competing alternatives and may be associated with reduced phonological or articulatory retrieval demands. Increased activation for word cues relative to phonemic cues in posterior dorsal supramarginal gyrus is of interest and consistent with the suggestion that the region may play an important role in the integration of semantic and phonological processes. These results provide an integral link between lesion studies and functional neuroimaging studies of language production in healthy speakers. Phonemic cuing of naming is argued to be an appropriate rehabilitation strategy for a variety of different patient groups as stages of semantic retrieval and phonological encoding are necessarily engaged during the task of overt naming (Best et al., 2013), thereby activating a network of regions that support these processes. The pattern of activity identified by the study in response to phonemic cues within an unimpaired system raises the possibility of using localized activity as a potential biomarker for interventions for word finding impairments after stroke.

Indeed, one approach has been to stimulate regions within the left-lateralized naming network using transcranial direct current stimulation (tDCS) during treatment paradigms (e.g., Baker et al., 2010, Fridriksson et al., 2011). In a recent fMRI study we demonstrated that anodal tDCS delivered to the LIFC during a cued picture naming study enhanced the behavioral and neural priming effect further in healthy older adults (Holland et al., 2011). Importantly, the degree of behavioral priming correlated with the magnitude of the neural priming effect in a regionally specific area within the LIFC. The ability to use localized neural responses as a possible target within a neural network for treatment interventions in patients with word finding difficulties would be an exciting development for neurorehabilitation.

## Acknowledgements

This study was supported by Wellcome funding (106161/Z/14/Z to JC) (203147/Z/16/Z to WCHN) and the Medical Research Council (G0701888). The funders had no participation in the design and results of this study.

